# Early Dynamics of Chronic Myeloid Leukemia on Nilotinib Predicts Deep Molecular Response

**DOI:** 10.1101/2021.02.17.431221

**Authors:** Yuji Okamoto, Mitsuhito Hirano, Kai Morino, Masashi K. Kajita, Shinji Nakaoka, Mayuko Tsuda, Kei-ji Sugimoto, Shigehisa Tamaki, Junichi Hisatake, Hisayuki Yokoyama, Tadahiko Igarashi, Atsushi Shinagawa, Takeaki Sugawara, Satoru Hara, Kazuhisa Fujikawa, Seiichi Shimizu, Toshiaki Yujiri, Hisashi Wakita, Kaichi Nishiwaki, Arinobu Tojo, Kazuyuki Aihara

## Abstract

Chronic myeloid leukemia (CML) is a myeloproliferative disorder caused by the *BCR-ABL1* tyrosine kinase^1,2^. *ABL1*-selective tyrosine kinase inhibitors (TKIs) including nilotinib have dramatically improved the prognosis of patients with CML^3–7^. The ultimate goal of CML treatment is likely to be TKI-free maintenance of deep molecular response (DMR), which is defined by quantitative measurement of *BCR-ABL1* transcripts on the international scale (IS)^8^, and durable DMR is a prerequisite to reach this goal^9^. Thus, an algorithm to enable the early prediction of DMR achievement on TKI therapy is quite valuable. Here, we show that our mathematical framework based on a clinical trial dataset^10^ can accurately predict the response to frontline nilotinib. We found that our simple dynamical model can predict nilotinib response by using two common laboratory findings (observation values): IS^11,12^ and white blood cell (WBC) count. Furthermore, our proposed method identified patients who failed to achieve DMR with high fidelity according to the data collected only at three initial time points during nilotinib therapy. Since our model relies on the general properties of TKI response, our framework would be applicable to CML patients who receive frontline nilotinib or other TKIs in clinical practice.

**Significance Statement:** Chronic myeloid leukemia (CML) is a myeloproliferative disorder caused by the BCR-ABL1 tyrosine kinase. The goal of this treatment is the sequential achievement of deep molecular response (DMR). Tyrosine kinase inhibitors (TKIs) are effective in the reduction because they inhibit CML cell proliferation. However, because of individual differences in the TKI efficacy, some patients are unable to achieve DMR, so that early prediction of DMR reachability is necessary for personalized medicine. By combining time series analysis and mathematical modeling, we developed a mathematical method that accurately predicts patients who do not achieve DMR in the early stages of TKI administration. Our prediction method gives a basis of effective personalized treatments for CML patients.

## Introduction

Chronic myeloid leukemia (CML) originates from a hematopoietic stem cell affected by a reciprocal translocation between chromosomes 9 and 22, which results in the formation of the *BCR-ABL1* fusion gene in the short derivative of chromosome 22 (Ph chromosome)^1^. In this rearrangement, latent ABL1 tyrosine kinase is constitutively activated by its oligomerization mediated by the N-terminal region of BCR and causes an unregulated expansion of myeloid cells, especially mature granulocytes^2^. CML is generally diagnosed during the stable chronic phase (CP). Since CP-CML cells are quite dependent on BCR-ABL1 for their growth and survival, selective ABL1 tyrosine kinase inhibitors (TKIs) have clinically demonstrated substantial reduction of *BCR-ABL1*-positive cells and restoration of normal hematopoiesis in the vast majority of CP-CML patients^3–7^.

At present, frontline TKIs for newly diagnosed CP-CML include imatinib^3^, dasatinib^4^, nilotinib^5–7^ and bosutinib^13^, among which the latter two are second generation TKIs with more potent activity than the former one. The actual therapeutic response to TKIs is estimated by quantitative polymerase chain reaction analysis of *BCR-ABL1* transcripts in peripheral blood and is represented by the ratio of *BCR-ABL1* mRNA to *ABL1* mRNA on the international scale (IS)^8, 11, 12^. The consensus optimal response to TKIs in CP-CML is major molecular response (MMR, IS ≤ 0.1%) within one year of therapy^14^, whereas a significant number of CP-CML patients achieve superior or deep molecular response (DMR; MR4.5, IS ≤ 0.0032%)^9, 15^, especially on frontline nilotinib or dasatinib^4–7^. Furthermore, a recent series of clinical trials on TKI discontinuation revealed that nearly a half of CP-CML patients maintained treatment-free remission (TFR) following more than two years of DMR on TKIs^16^. Thus, to ultimately reach TFR, the practical target of CML treatment is to achieve DMR as early as possible after frontline TKI therapy, and the switch to another TKI may be a therapeutic option when DMR is not likely to be obtained. Thus, an algorithm to enable the early prediction of DMR achievement on TKI therapy would be quite valuable.

In this paper, we propose a mathematical framework that can predict the efficacy of nilotinib in each CML patient only from short time-series data. We constructed a simple classification method that correctly estimated whether a CML patient reached DMR or not within 24 months after nilotinib administration. The key to success is to focus on the dynamical behavior of CML cells.

## Results

The dataset analyzed in this study was obtained in a phase II clinical trial named the N-road study^10^ that aimed to examine the safety and efficacy of nilotinib for newly-diagnosed CML patients. Complete blood counts (CBCs) and IS were measured in a single certified laboratory center every 3 months for 2 years. In this study, we analyzed the data from the eligible 32 patients. Hereafter, among the eligible patients, we defined 18 patients who have reached DMR by 24 months as DMR patients and remaining 14 patients as non-DMR patients (Fig. S1).

Recently, K. Sasaki *et al.* reported optimal IS values that predict the highest probability of reaching a sustained DMR at specific time points between 3 and 12 months^17^. However, to the best of our knowledge, there are neither scoring systems nor algorithms for early prediction of DMR achievement in CML patients under TKI therapy. Therefore, we examined whether the current scoring systems and guidelines in CML clinical practice could be adapted to predict DMR. The European LeukemiaNet (ELN) guideline in 2013^14^ recommends the EUTOS prognostic score for prediction of complete cytogenetic response (CCyR) and progression-free survival (PFS) on frontline imatinib^18^. In the N-road study dataset, the EUTOS score did not precisely discriminate DMR from non-DMR patients (Fig. 1a). We tested another prognostic score, EUTOS long-term survival (ELTS) score, which is newly adopted in ELN guideline 2020^19^. The ELTS score was developed for prediction of long-term survival considering leukemia-related death. As well as the EUTOS sore, the ELTS score also did not distinguish between DMR and non-DMR patients in the N-road study (Fig. 1b). In addition, the definition of response to frontline TKIs employed in the European LeukemiaNet (ELN)^14,19^ required at least 12-month observation to discriminate the two populations in the same cohort (Fig. 1c-e). Then, we sought for key features to construct a better method of early DMR prediction.

**Figure 1.**
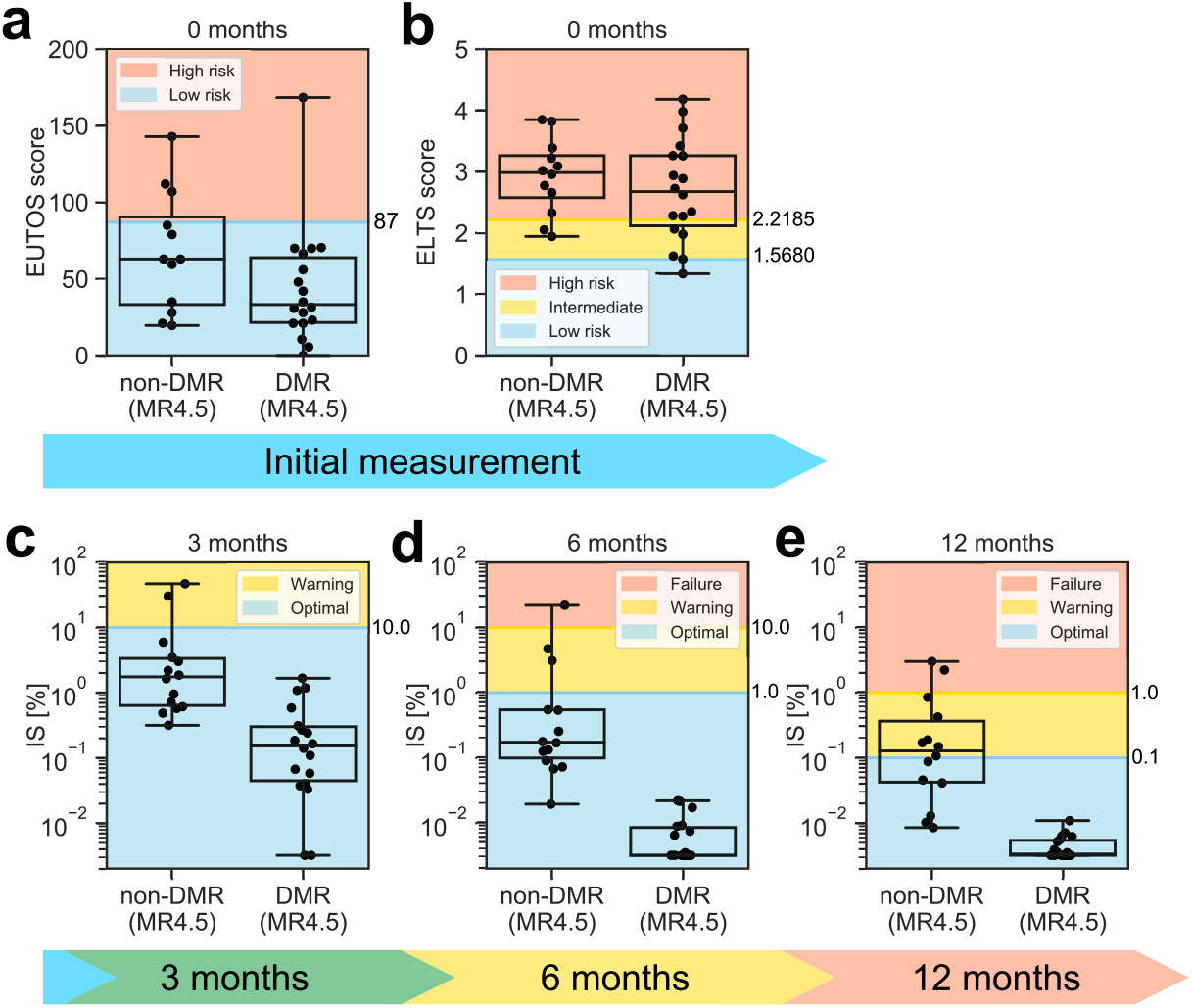
Several criteria of the current scoring systems and guidelines are not suitable for DMR classification. Each dot indicates each patient value. The distributions of these values are presented as box-plots. The top and bottom bars indicate the maximum and minimum values, respectively. The top, center, and bottom lines of the boxes are the lower quartiles, medians, and upper quartiles, respectively. **a**, We calculated the EUTOS score for each patient within the N-road dataset. The patient is considered “High risk” if the EUTOS score of a CML patient is higher than 87 and “Low risk” otherwise^18^. We divided patients into two groups: a DMR patient set and a non-DMR patient set. The EUTOS score did not predict non-DMR patients, because most of the non-DMR patients were predicted as “Low risk”. **b**, The ELTS score for each patient was also calculated within the N-road dataset. A CML patient is considered as “High risk” if the patient’s ELTS score is higher than 2.2185, “Low risk” if the score is 1.5680 or lower, and “Intermediate risk” otherwise^19^. The ELTS score predicted most patients as “High risk” or “Intermediate risk”. These predicted results imply that the ELTS score did not correctly select non-DMR patients. **c-e**, According to the ELN guideline^14^, we plotted patient IS and separated them into 3 groups: “Optimal”, “Warning”, and “Failure” for the data at (**c**) 3, (**d**) 6, and (**e**) 12 months after the initiation of TKI administration. In **c** and **d**, many non-DMR patients were predicted as “Optimal”, indicating the failure of these predictions. Their performance was evaluated in terms of accuracy, sensitivity, and specificity. Results are summarized in Fig. 4. Detailed information can be found in the Materials and Methods.

Potential key features should be found in the dynamics of *BCR-ABL1*-positive cells (called CML cells in the following) in CML patients on TKIs because mathematical studies on CML^20–25^ explained the characteristic dynamics of *BCR-ABL1*-positive cells in CML patients on TKIs and successfully predicted relapse. Particularly, M. Horn *et al.* showed that relapse was predictable based on residual leukemic stem cells and reduction slopes of IS^23^. However, for the purpose of DMR prediction from short-term measurements, most models are too complex to obtain suitable parameter values for each patient. Numerous approaches have been proposed to overcome these data-shortage problems such as high-dimensional data in general^26^ and prostate cancer in specific^27,28^. In this research, we overcome this problem by focusing on two important key features on the dynamics of BCR-ABL1-positive cells in CML patients and by proposing an adequately-simple mathematical model, which can reproduce the two key features from a small dataset.

Two key features for early DMR prediction in CML patients are the decreasing rate and the convergence value of IS time-series data (Fig. 2). Both early and late dynamics of IS can be clearly represented by these two features (Fig. 2a and b). Additionally, the distributions of these two features in DMR and non-DMR patients seem to be distinguishable (Fig. 2c and d). Thus, we can discriminate DMR patients from non-DMR patients, if we accurately estimate these two features from short time series of IS values. Hereafter, we discuss a mathematical way to estimate these two features.

**Figure 2.**
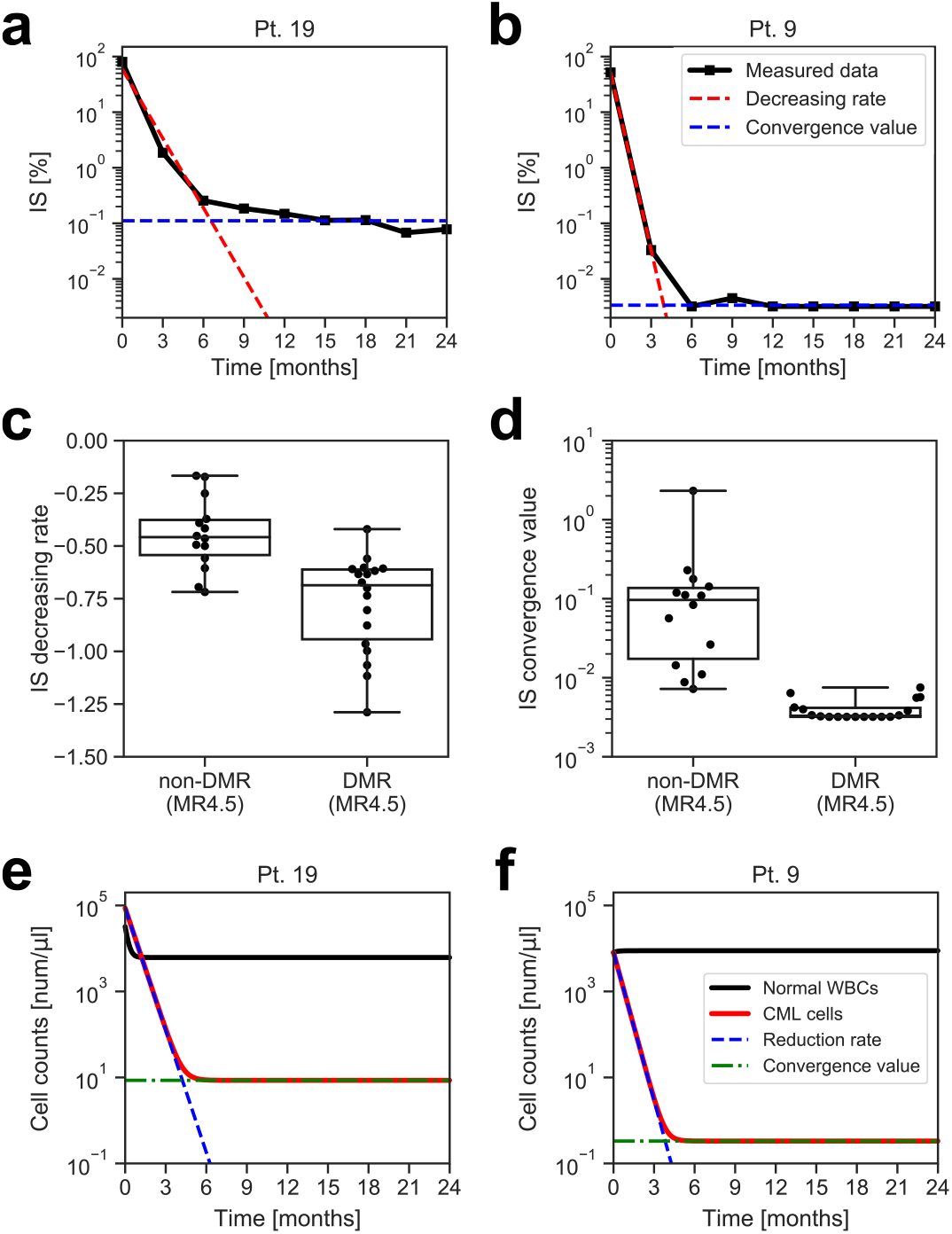
Key features for DMR prediction: the decreasing rate and convergence value of IS, and the reduction rate and convergence value of CML cells. **a**, **b,** The time series of IS are plotted in **a** for a non-DMR patient (patient # 19) and in **b** for a DMR patient (patient # 9). The decreasing rate and convergence value of IS are two key features for DMR prediction. A detailed information of the parameter estimation can be found in the Materials and Methods. **c,** The distribution of the IS decreasing rate for non-DMR and that for DMR patients were clearly different. **d,** The distribution of the IS convergence value for non-DMR and that for DMR patients were clearly distinct. **e, f,** We show the estimated dynamics of normal WBCs and CML cells counts in **e** for a non-DMR patient (patient # 19) and in **f** for a DMR patient (patient # 9). These figures clearly indicate that the reduction rate and convergence value of CML cells play a key role in the distinction between DMR and non-DMR patients.

First, we propose a mathematical model, which describes the time evolution of the white blood cell (WBC) count in CML patients. Our model includes two variables: the normal WBC (namely *BCR-ABL1*-negative) count and CML cell (namely *BCR-ABL1*-positive) count. Furthermore, each variable has two parameters: the recovery rate and the convergence value for the normal WBC count, and the reduction rate and the convergence value for the CML cell count. Since the both cell counts are mathematically calculated by the measurement of IS and the total WBC count in our model (Eqs. 2-3 in Materials and Methods), we can estimate the variables and parameters (Fig. 2e-f). The first feature, the decreasing rate of IS, corresponds to the reduction rate of the CML cells. The second feature, the convergence value of IS, corresponds to the convergence value of the CML cells. Our mathematical model sufficiently approximates the time series of IS for various CML patients (Fig. 3a). These findings indicate the validity of our proposed model (Materials and Methods, and Fig. S2).

**Figure 3.**
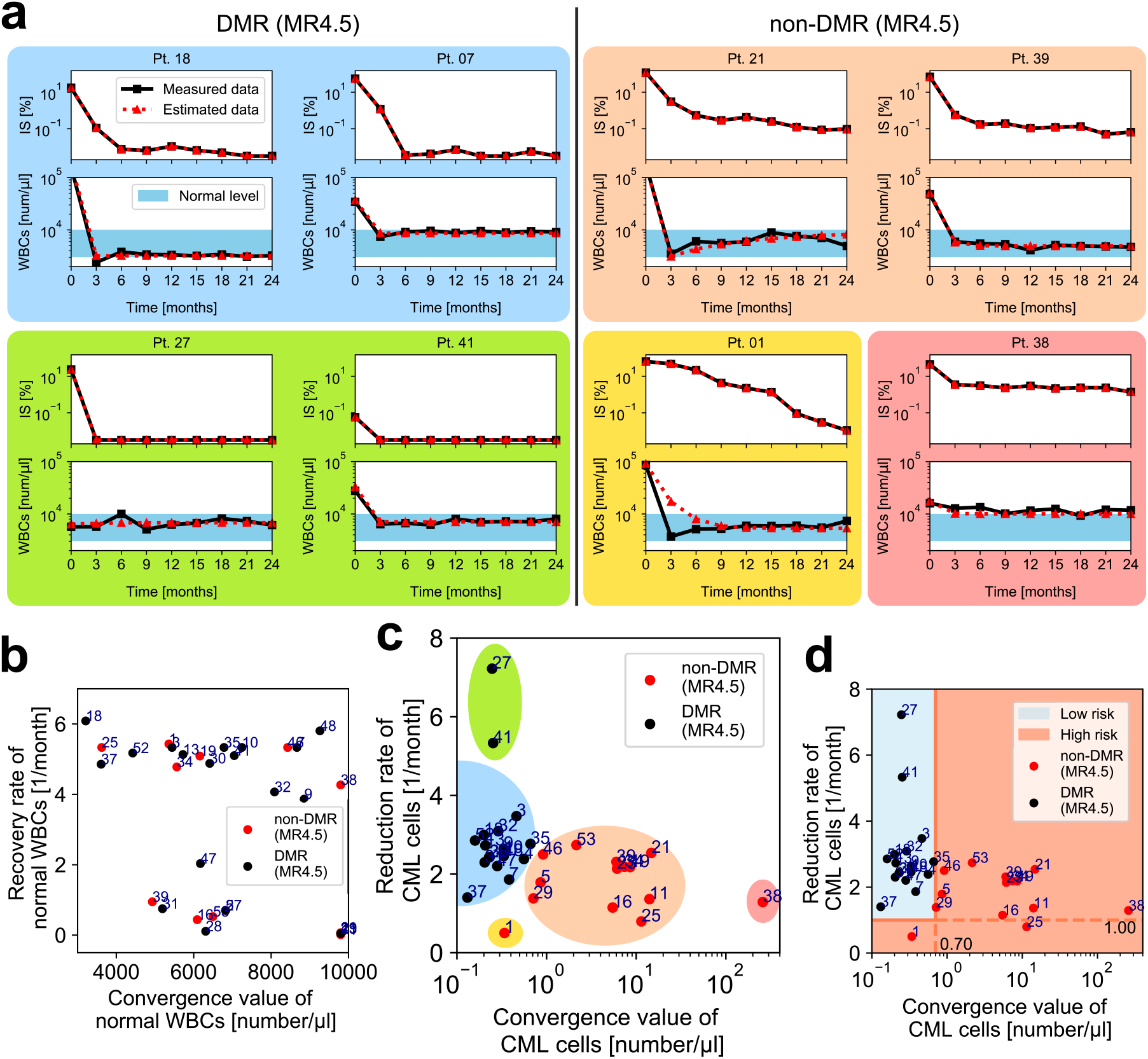
DMR and non-DMR patients were separated on the plane of the reduction rate and convergence value of CML cells. **a**, The estimated time series of CML dynamics (red dashed lines) sufficiently approximated the measured data for IS [%] and for WBC counts [number / μl] (black solid lines). **b**, The distribution of the convergence value and recovery rate [1 / month] of normal WBCs did not show a correlation in terms of nilotinib response. The number indicated on each dot is patient #. **c**, The distribution of the convergence value and reduction rate of CML cells was apparently classified into several subsets: a set including most DMR patients (the blue region), non-DMR patients (the orange region), and outlier patients (other color regions). **d**, By choosing optimal threshold values, the distribution of DMR patients shown in **c** was perfectly separated from that of non-DMR patients. Notably, this result was obtained by using all patient data points.

Second, we discuss how we deal with estimated parameter values to classify the TKI response (Fig. 3b-d). DMR and non-DMR patients were clearly separated on a parameter plane of the CML cell reduction rate and convergence value (Fig. 3c). By setting appropriate threshold values determined with all the eligible patient data points, we positively discriminated between DMR and non-DMR patients in the analyzed N-road dataset with 100 % accuracy (Fig. 3d). On the other hand, another set of parameters did not clearly separate patients (Fig. 3b). We note that, in Fig. 3, all the time points of all patients data were used to estimate our model parameters. However, the parameters can be estimated based only on three time points because of our model’s simplicity. To classify patients into DMR or non-DMR based on the estimated parameters, the threshold values still need to be determined.

Third, we discuss an approach to determine the threshold values in a CML patient by utilizing information from other patients. We employed the leave-one-out cross-validation. By applying the aforementioned framework to data from other patients, we obtained a scatter diagram and optimized the threshold values under a certain criterion. In Fig. 4, we summarize the performance of DMR predictions. Our proposed approach, which uses data only at 3 initial time points (at 0, 3, and 6 months), reached the performance of the ELN definition at 12 months of therapy. Furthermore, our method enhanced specificity compared to the EUTOS scoring system and ELN guidelines. As for ELTS scoring system, it estimated most patients as “Intermediate risk” or “High risk”. By assuming that these classifications correspond to non-DMR, we consider the ELTS scoring system to be difficult to detect non-DMR patients. Thus, our algorithm has the high potential to predict non-DMR patients on frontline nilotinib, and our method may improve the current CML treatment.

**Figure 4.**
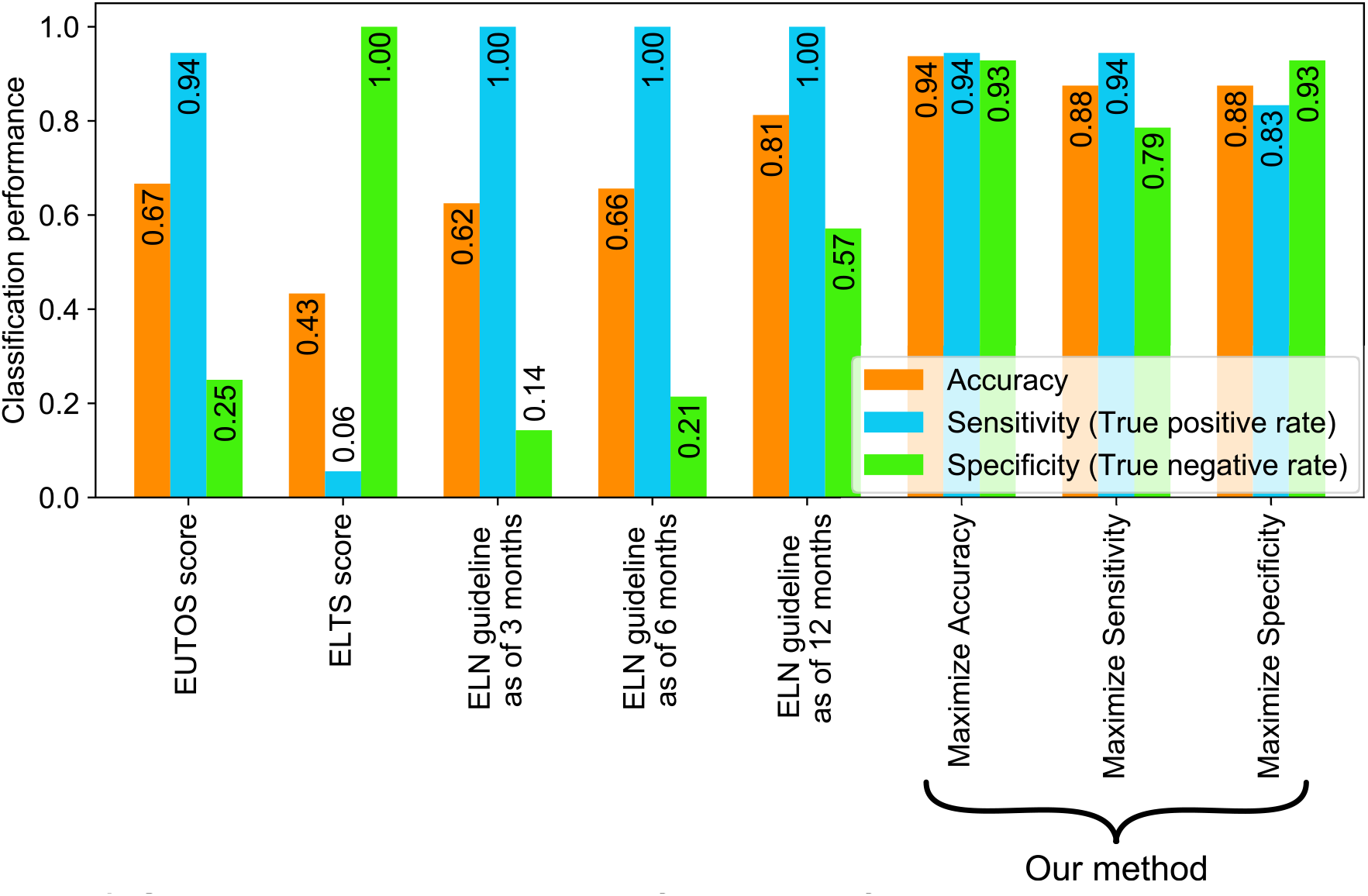
Our prediction method achieved favorable performance in comparison with the criteria of the current scoring systems and guidelines. We summarize the performance for accuracy (orange bars), sensitivity (blue bars), and specificity (green bars) of the EUTOS score, the ELTS score, the ELN guideline for the cases of 3, 6, 12 months, and our proposed method (presented with the three rightmost labels). In the current study, “positive” and “negative” indicate DMR and non-DMR patients, respectively. In the cases of the EUTOS scores, the ELTS scores, and the ELN guidelines, we decided to estimate a target patient as “positive”, if the patient was classified as “Low risk” or “Optimal”. Under this estimation, we obtained accuracy, sensitivity (the true positive rate), and specificity (the true negative rate) for each case. In our setting, the prediction of non-DMR patients is highly important, because the administrated TKI should be promptly changed for them. In this sense, our method was superior to the EUTOS score and the ELN guideline because our proposed method showed higher specificity than the scoring system and guideline. The ELTS score showed higher specificity than our proposed method because most patients were estimated as negative. Thus, the ELTS score cannot distinguish between non-DMR and DMR patients, which was also reflected in the low sensitivity. Notably, each of our three results was optimized in terms of accuracy, sensitivity, and specificity through the training period.

## Discussion

Finally, we discuss whether the presented mathematical model is applicable to other frontline TKIs including imatinib and dasatinib. The BCR-ABL1 transcript dynamics under imatinib treatment^21, 24^ was similar to the IS dynamics under nilotinib treatment analyzed in our study. Although further investigations are needed, we expect that our model may be applicable to imatinib treatment. Dasatinib induces a rapid and significant mobilization of normal lymphocytes and monocytes in peripheral blood with a peak around 2 hours after oral intake and their cell counts reflect its plasma concentration^29^. This means that the WBC count, one of the two key model features, is likely to be affected by dasatinib irrespective of the dynamics of CML cells. However, the time scale of this phenomenon is much shorter than that of our mathematical model. Due to the timescale separation, the fast dynamics induced by dasatinib may not affect the slow dynamics of WBCs and CML cells. If so, the present model is also valid for dasatinib.

In conclusion, our mathematical method accurately predicted the possibility to achieve DMR in a retrospective cohort of CML patients treated with frontline nilotinib. This outcome is due to the suitable degree of simplicity of our model: it is simple enough to operate with short-term time series but also complex enough to distinguish between DMR and non-DMR patients. The ultimate goal of CML treatment is likely to be TKI-free maintenance of DMR, and the achievement of durable DMR is a prerequisite for this goal^30^. Furthermore, timely prediction of nilotinib failure to achieve DMR will enable the consideration of other therapeutic approaches, which will allow sustained DMR leading to TFR. Therefore, the present study contributes to the progress of mathematical applications in clinical settings.

## Materials and Methods

### Definition of deep molecular response (DMR)

In this study, we employed DMR as the criterion for effective drug response in CML patients. There are multiple definitions of DMR such as MR4.0 (IS ≤ 0.01%) and MR4.5 (IS ≤ 0.0032%)^8, 9, 14, 15, 19^. If a patient’s IS becomes lower than one of the above threshold values at a certain time point, it is considered that the patient reaches the DMR corresponding to the employed threshold value. There is a stricter criterion designated as complete molecular response (CMR). A patient is considered to reach CMR, if his/her expression of the BCR-ABL1 gene is not measurable at two consecutive time points^8, 31^. In this paper, we designated patients as CMR who reached MR4.5 at both 21 and 24 months. Because lower IS implies a more favorable situation for CML patients, CMR is considered to be a better criterion. However, the measurement of low BCR-ABL1 expression is technically challenging, i.e., non-negligible noise/error is often contained in such measurements. Therefore, we employed MR4.5 for this analysis. In Fig. S3-5, for MR4.0, MR4.5, and CMR, we compared the numerical results corresponding to Fig. 1, Fig. 3b-d, and Fig. 4.

Here, we note a more detailed definition of DMR patients. We designated patients whose IS decreased below 0.0032% at a certain time point as DMR patients. However, in a subset of these DMR patients, their IS fluctuated and/or again increased above 0.0032%. In this study, we treated these patients as DMR patients. In further investigation, this point may be reconsidered, if more detailed clinical information becomes available. One of the reasons for IS reoccurrence may be the acquisition of drug resistance. However, we cannot distinguish patients who acquired drug resistance based on the available information, because observations over a longer time span are needed.

### Detailed information of the N-road trial

We used the dataset from the N-road trial^10^, which was a phase `II`, open-label, multi-center clinical study for newly diagnosed CP-CML patients. This study was organized based on previous trials on nilotinib^5–7^. The CP-CML diagnosis had to be made by chromosome banding or fluorescent in situ hybridization (FISH) or reverse transcription polymerase chain reaction (RT-PCR). The N-road aimed to investigate the safety and efficacy of nilotinib in newly diagnosed CP-CML patients based on the early achievement of CMR. The primary endpoint was the CMR (defined as MR4.5 in the N-road study) achievement rate within 24 months. This trial included 53 patients and was conducted from August 2012 until July 2017. Nilotinib 300 mg twice daily was administered to patients for 24 months. If patients did not reach the criteria for optimal response defined by the N-road study referring to the ELN 2013 definition^14^ at any time point, the administration of nilotinib up to 400 mg twice daily as second-line therapy was allowed.

Notably, our mathematical framework requires only IS and WBC counts. Other laboratory findings were also obtained but were not used in this paper. In addition, in this study, we analyzed the data from the eligible 32 patients; the remaining 21 patients were ineligible for analysis due to missing data of one or more CBC and/or BCR-ABL1 IS measurements. Furthermore, in this paper, moving to the second line therapy or not was not thematized.

This trial was approved by the institutional review boards of the participating institutions and was conducted in accordance with the ethical guidelines for medical and health research involving human subjects. This study also adhered to the ethical principles of the Declaration of Helsinki. All patients provided written informed consent to participate in the study. This N-road study is registered in University hospital Medical Information Network (UMIN) system, and its ID is UMIN000008565^10^.

### Details of the EUTOS score, the ELTS score, and the criteria of the current guideline

Here, we introduce details of the EUTOS score^18^, the ELTS score^19^, and the ELN guidelines from 2013^14^ and 2020^19^ to be related to our study. Both the EUTOS score^18^ and the ELTS score^19^ are the prognostic scores to estimate the survival risk at baseline (i.e., before the TKI administration). The EUTOS score^18^ was proposed as a score to predict CCyR and subsequently PFS. The EUTOS score is calculated by the basophil ratio and splenomegaly. If the EUTOS score of a patient is higher than 87, the patient is designated as being in a “High risk” situation. The ELTS score was recently adopted in the ELN guideline 2020^19^ and proposed for long-term survival considering leukemia-related death^19^. At baseline, the ELTS score is calculated by the age in completed years, the spleen size below costal margin, the percentage of blasts in peripheral blood, and the platelet count. A CML patient is considered as “High risk” if the patient’s ELTS score is higher than 2.2185, “Low risk” if the score is lower than 1.5680, and “Intermediate risk” otherwise.

In the ELN guideline 2013^14^, the condition of a patient is classified into three stages according to the relationship among IS, chromosome banding analysis (CBA) from bone marrow cells, FISH of blood interphase cell nuclei, and duration since the beginning of TKI administration: The three stages are “Failure”, “Warning”, and “Optimal” starting from the worst. At the 3-month time point, the threshold value between “Warning” and “Optimal” is set at 10% of IS. At the 6-month time point, the threshold value between “Failure” and “Warning” is set at 10%, and that between “Warning” and “Optimal” is at 1%. At the 12-month time point, these threshold values are set at 1% and 0.1%, respectively. The above information is illustrated in Fig. 1 of the main text.

We should note that the ELN guideline 2020^19^ changed the criteria of the risk stage. However, the changes did not affect the results obtained from our dataset. Here, we explain the change related to our analysis and why it did not affect our result. One of the changes related to our analysis is the “Failure” stage criteria at the 3-month time point. The criteria of the guideline in 2013^14^ do not explicitly refer to IS values, but those in 2020^19^ refer to IS values. According to the guideline in 2020, a patient is confirmed as the “Failure” stage if the IS value satisfies the following both conditions: the IS value is higher than 10% at the 3 months; the same IS level is confirmed again before the 6 months. The N-road dataset has IS values every 3 months. Therefore, no patient satisfied the second condition. Thus, the changes in the ELN guideline 2020 did not affect our results.

We tested the performance of the EUTOS score, the ELTS score, and the ELN guideline for DMR prediction in our dataset as shown in Fig. 4 of the main text. It should be noted that the EUTOS score, the ELTS score, and the guideline were not proposed for DMR prediction. In the cases of the EUTOS score and the guideline with 3 months and 6 months (labeled as “ELN guideline as of 3 months” and “ELN guideline as of 6 months”), the results indicated high “sensitivity” and low “specificity”; therefore, the prediction of non-DMR patients based on them is very difficult. On the contrary, for the ELN guideline as of 12 months, specificity was higher at 57%. Thus, for accurate prediction of non-DMR patients based on the guideline 12 months duration is necessary. The ELTS score has higher “specificity” than the ELN guideline as of 12 months. However, the “sensitivity” and “accuracy” are significantly low. This result indicates that the ELTS score is not suitable for the prediction of DMR patients (see Fig. S3-5 for the performance of the EUTOS score, the ELTS score, and the guideline in the cases of MR4.0, MR4.5, and CMR). The performance of the current guidelines, excluding the EUTOS score and the ELTS score, was tested in 32 patients. Since spleen size data, which are required to calculate the EUTOS and the ELTS scores, were not available for two patients, the EUTOS and ELTS scores were evaluated in the remaining 30 patients.

### Dynamical model of CML cells

Here, we introduce the details of our dynamical model. We modeled the dynamics within peripheral blood of CML patients. Administration of an effective TKI results in the reduction of CML cells (BCR-ABL1-positive WBCs) and the increase of normal WBCs (BCR-ABL1-negative WBCs) in peripheral blood. In this study, we assumed that the number of CML cells and that of normal WBCs exponentially change with time. We also assumed that these numbers asymptotically converge to constant values. Then, the time series of these numbers can be described as a couple of linear differential equations as follows:

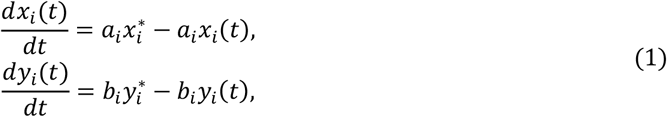

where *x_i_*(*t*) and *y_i_*(*t*) represent the number of normal WBCs and that of CML cells for the *i*th patient at time *t*, respectively. Their convergence values are denoted by 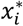 and 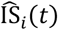. The coefficients *a_i_* and *b_t_* represent the recovery rate of normal WBCs and the reduction rate of CML cells for the *i*th patient, respectively. Examples of the dynamics generated by the model are shown in Fig. 2e-f.

### Relationship between measured values and CML dynamics

We explain the relationship between our model and measurements from CML patients. We can convert two measurement variables, the WBC count and IS, into two state variables in our model, *x_i_*(*t*) and *y_i_*(*t*), via the following procedure. The WBC count clearly agrees with the sum of *x_i_*(*t*) and *y_i_*(*t*). Then, the WBC count for the *i*th patient at time t, denoted by WBC¿(t), is described as follows:

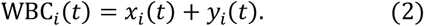

The definition of IS is the ratio of the BCR-ABL1 gene (mRNA) expression to the ABL1 gene (mRNA) expression: The BCR-ABL1 gene is expressed in CML cells, whereas the ABL1 gene is expressed in both CML cells and normal WBCs. Here, we assume that gene expression is proportional to the number of cells. We denote the ratio of gene expression in normal WBCs to that in CML cells by *c*. Thus, IS for the *i*th patient at time *t*, denoted by IS_*i*_(*t*)[%], is described as follows:

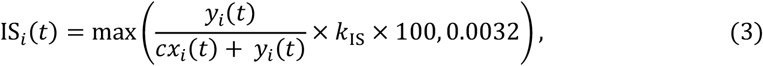

where *k*_IS_ is the conversion factor of IS, and our dataset of the N-road adopts k_IS_ = 1.2. Because the detection limit of IS measurements in this dataset is 0.0032%, we set the lower bound as 0.0032.

These equations (2) and (3) can transform the WBC count WBC_*i*_(*t*) and IS IS_*i*_(*t*) into *x_i_*(*t*) and *y_i_*(*t*), respectively. However, the constant ratio c is generally unknown. Thus, we estimated the ratio c and the parameters of the CML dynamical model 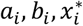, and 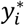 from the measurements as described in the following section.

### Fitting IS time series data with piecewise functions

In this part, we explain the method to estimate IS dynamics parameters shown in Fig. 2c and d. To obtain the decreasing rate and convergence value of IS, we defined a piecewise function with two domains as follows:

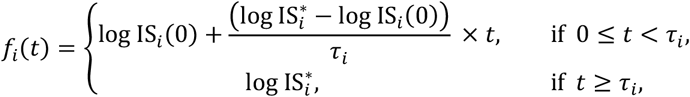

where 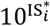 is the IS convergence value, 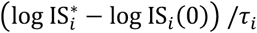 is the IS decreasing rate of the *i*th patient, and *τ_i_* [month] is a boundary between the two domains. Then we solve the following optimization problem:

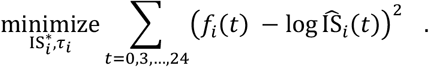

We used the Adam optimizer of the TensorFlow machine learning library to solve the problem.

### Parameter estimation of the CML dynamical model

Here, we introduce the optimization method to estimate the parameters of equations (1), (2), and (3). The measurement value of the WBC count and that of IS for the *i*th patient at the time *t* are denoted by 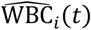 and 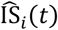, respectively. Then, the optimization problem is described as follows:

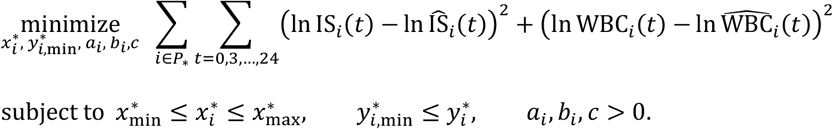

As for the fitting IS data with the piecewise function, we used the Adam optimizer to solve this optimization problem. We employed the method of Lagrange multipliers, and the constraint is

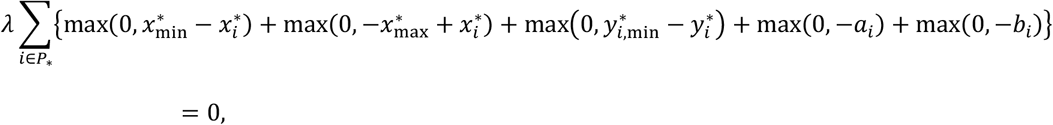

where *λ* = 100. We set the lower bound 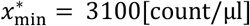 and the upper bound 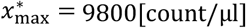 in the constraint condition to maintain the convergence values of normal WBCs within the normal levels of healthy humans. Furthermore, we set the lower bound 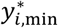 in the constraint condition to keep IS_*i*_(*t*) beyond the detection limit of IS, i.e., 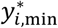 is obtained by solving 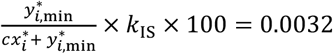. A set of patient indices is denoted by *P*\*. In Fig. 3, the parameters are estimated by utilizing all patient data, i.e., *P\** contains all patients (*i* = 1,…,32). In Fig. 4, we checked how our proposed method outperformed the existing guidelines in terms of DMR patient prediction using short time series. For this sake, we did leave-one-out crossvalidation, i.e., *P\** contained 31 patients used for the training dataset, and the remaining one patient was the prediction-target patient. We repeated this data splitting 32 times employing subsequently all 32 patients as the prediction target patient.

### Parameter Estimation for the Prediction-Target Patient

Here, we introduce an approach to estimate parameters of the prediction-target patient only from three data points. As we have discussed in Fig. 3b-d, the classification is performed on the plane of the CML cell convergence value 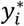 and its reduction rate *b_i_*. Thus, we introduce a method to estimate the two parameters from three data points, which are specifically denoted by 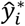 and 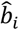. Notably, the gene expression ratio parameter *c* is already estimated in the previous training step. Hereafter, we assume that all ISs of the three data points are greater than the detection limit. Otherwise, clearly according to the definition of DMR, the patient must be a DMR patient. Under this assumption, from equations (2) and (3), the estimated number of CML cells at time t, denoted by 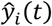, is obtained as follows:

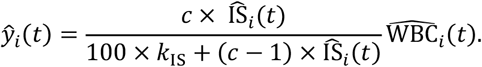

Then, the estimated convergence value 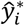 and the reduction rate 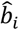 are obtained as follows:

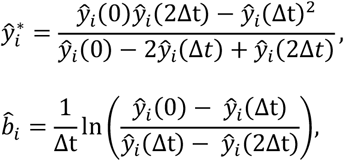

where Δ*t* represents the interval of measurement points. In our dataset, the N-road, the data are measured every 3 months, i.e., Δt = 3[month], meaning that we utilized the data 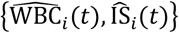 for t = 0,3, and 6 [months]. Based on the estimated parameters 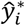 and 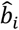, our model predicts DMR or non-DMR for the target patient. Note that, according to our DMR definition, we regard a patient as DMR, if the patient’s IS value satisfies the DMR definition (IS ≤ 0.0032% for MR4.5) for t = 0,3, or 6 [months].

### DMR Prediction Criteria

We introduce an approach to determine the criteria for DMR prediction. As presented in Fig. 3d, DMR patients should show large enough *b_i_* and small enough 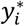. To estimate the parameters 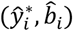, we define a parameter region 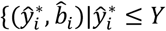 and 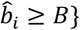 for the criteria. If the estimated parameters 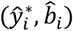 are included in this region, the patient is predicted to be a DMR patient. These threshold values *Y* and *B* are determined by using the training data, i.e., patients whose ID are included in a set *P*\*. We explored the optimal *Y* and *B* that maximize the accuracy, sensitivity, or specificity. We used the exploration intervals 0.1 and 0.05 for *Y* and *B*, respectively. These indices are the common concept in the classification context as follows. The accuracy is the ratio of patients who are successfully predicted. The sensitivity, the true positive rate, is the ratio of DMR patients who are successfully predicted. The specificity, the true negative rate, is the ratio of non-DMR patients who are successfully predicted. As described above, if a patient satisfies the DMR definition within its initial three time points, the patient is considered to be a DMR patient. In Fig. 4 and Fig. S5, the numbers of such patients were 15 for MR4.0, 10 for MR4.5, and 2 for CMR. All these patients were estimated to be “Optimal” in the case of the ELN guidelines^14, 19^. For MR4.0 and MR4.5, only one patient of them was estimated to be “High risk” in the case of the EUTOS score, and the remains were estimated to be “Low risk”. For CMR, all the patients were estimated to be “Low risk” in the case of the EUTOS score. In the case of the ELTS score, only one of the patients for MR4.0 and MR4.5 was estimated to be “Low risk” and the remains were estimated to be “High risk” or “Intermediate risk”, and all of the patients for CMR were estimated to be “Intermediate risk”. Here, for each accuracy, sensitivity, and specificity, we remark how to choose the optimal threshold (*Y,B*). In the case of maximizing accuracy, we obtained the multiple possible sets of optimized (*Y,B*) that maximize accuracy. Then, from the possible sets, we selected the set (*Y,B*) closest to the average values of the possible sets. In the case of maximizing sensitivity, we obtained the multiple possible sets (*Y,B*) which achieve the largest sensitivity. However, under this condition of maximizing sensitivity, we can freely magnify the parameter region corresponding to DMR. To keep the area of the region reasonable, from the above possible sets, we again selected parameter sets that achieve the maximal true negative rate. Finally, we selected the largest *Y* and the smallest *B* among the sets that had passed the both selectins as the optimal set. In the case of maximizing specificity, similar to the case of sensitivity, we obtained the multiple possible sets (*Y,B*) which achieve the largest specificity. Then, from the possible sets, we again selected parameter sets that achieve the maximal true positive rate. Finally, we selected the smallest *Y* and the largest *B* among them as the optimal set.

Through this framework, by maximizing the accuracy, sensitivity, or specificity, we obtained three prediction results as shown in Fig. 4 (for MR4.5) and Fig. S5 (for MR4.0, MR4.5, and CMR). Our results with specificity maximization provided good performance especially in terms of specificity. Thus, this method can successfully predict non-DMR patients from short-term measurements. In future studies analyzing a larger dataset, we plan to employ machine learning methods for the optimization of our framework. The prediction of characteristic features of CML patients may also represent a future research avenue. In Fig. 3c, one can find that some patients were outliers, i.e., they were separated from most DMR and most non-DMR patients. Further clinical analysis on these outliers may provide new characteristic features of CML patients. As an example of related works, M. Horn *et al.* showed that patient classification on parameter space predicted relapse^22^. Thus, prediction of relapse may be another possible application of our mathematical framework.

### Disclosure of institutional review board approval

This research has been approved by the Institutional Review Boards at The University of Tokyo, at The Institute of Medical Science, The University of Tokyo (IMSUT), at Kyushu University, at University of Fukui, and at Hokkaido University.

## Acknowledgments

This work was partially supported by JSPS KAKENHI Grant Number JP15H05707 (to K. M., M. K. K., S. N., and K. A.), JST PRESTO Grant Number JPMJPR16E9 (to S.N.) and JST Moonshot R&D Grant Number JPMJMS2021 (to.K.A.). The N-road study was funded by Novartis Pharma K. K.

## Notes

### Competing Interest Statement

K. M. reports a grant from NEC outside the submitted work. S. N. reports a grant from Morinaga Milk Industry Co., Ltd. outside the submitted work. K. N. received a grant for this work from Novartis Pharma K.K. and reports grants from Zenyaku Kogyo Company, Ltd., Asahi Kasei Pharma, and Taiho Pharmaceutical Co., Ltd.; grants and personal fees from Chugai Pharmaceutical, Kyowa Hakko Kirin Co., Ltd., Nippon Shinyaku Co., Ltd., Mochida Pharmaceutical Co., Ltd., Ono Pharmaceutical Co., Ltd., Takeda Pharmaceutical Co., Ltd., and Sumitomo Dainippon Pharma Co., Ltd.; and personal fees from Pfizer, Otsuka Pharmaceutical Co., Ltd., Janssen Pharmaceutical K.K., Eisai Co., Ltd., and Celgene K.K. outside of the submitted work. H. Y. reports grants from Celgene K.K. and Astellas Pharma Inc. outside the submitted work. A. T. reports personal fees from Sysmex, Otsuka Pharmaceutical Co., Ltd., Bristol-Myers Squib, and Takeda Pharmaceutical Co., Ltd., Daiichi-Sankyo and grants and personal fees from Pfizer, and Chugai Pharmaceutical outside the submitted work. K. A. reports a grant and personal fees from KKE, a grant and personal fees from NEC, a grant from Sysmex, a personal fee from Novo Nordisk Japan, and a grant and personal fees from Toyota Central R&D Labs. outside the submitted work. Y.O. had worked at Sysmex Co., but the entire results of this paper are totally independent from his past job in Sysmex Co.. The other authors have nothing to disclose.

## References

1. E. Shtivelman, B. Lifshitz, R. P. Gale, E. Canaani. Fused transcript of abl and bcr genes in chronic myelogenous leukaemia. Nature. 315, 550–554 (1985).

2. J. V. Melo, D. J. Barnes. Chronic myeloid leukaemia as a model of disease evolution in human cancer. Nat. Rev. Cancer. 7, 441–453 (2007).

3. B. J. Druker, F. Guilhot, S. G. O’Brien, I. Gathmann, H. Kantarjian, et al., Five-year followup of patients receiving imatinib for chronic myeloid leukemia. N. Engl. J. Med. 355, 2408–2417 (2006).

4. M. Talpaz, N. P. Shah, H. Kantarjian, N. Donato, J. Nicoll, et al., Dasatinib in imatinib-resistant Philadelphia chromosome-positive leukemias. N. Engl. J. Med. 354, 2531–2541 (2006).

5. G. Saglio, D. W. Kim, S. Issaragrisil, P. Le Coutre, G. Etienne, et al., Nilotinib versus imatinib for newly diagnosed chronic myeloid leukemia. N. Engl. J. Med. 362, 2251–2259 (2010).

6. H. Kantarjian, F. Giles, L. Wunderle, K. Bhalla, S. O’Brien, et al., Nilotinib in imatinib-resistant CML and Philadelphia chromosome-positive ALL. N. Engl. J. Med. 354, 2542–2551 (2006).

7. H. M. Kantarjian, F. Giles, N. Gattermann, K. Bhalla, G. Alimena, et al., Nilotinib (formerly AMN107), a highly selective BCR-ABL tyrosine kinase inhibitor, is effective in patients with Philadelphia chromosome-positive chronic myelogenous leukemia in chronic phase following imatinib resistance and intolerance. Blood. 110, 3540–3546 (2007).

8. N. C. Cross, H. E. White, M. C. Müller, G. Saglio, A. Hochhaus. Standardized definitions of molecular response in chronic myeloid leukemia. Leukemia. 26, 2172–2175 (2012).

9. R. Hehlmann, M. C. Müller, M. Lauseker, B. Hanfstein, A. Fabarius et al., Deep molecular response is reached by the majority of patients treated with imatinib, predicts survival, and is achieved more quickly by optimized high-dose imatinib: results from the randomized CML-study IV. J. Clin. Oncol. 32, 415–423 (2014).

10. K. Nishiwaki, K. J. Sugimoto, S. Tamaki, J. Hisatake, H. Yokoyama, et al., Optimal treatment strategy with nilotinib for patients with newly diagnosed chronic-phase chronic myeloid leukemia based on early achievement of deep molecular response (MR^4.5^): The phase 2, multicenter N-Road study. Cancer Med. 9, 3742–3751 (2020).

11. T. Hughes, M. Deininger, A. Hochhaus, S. Branford, J. Radich, et al., Monitoring CML patients responding to treatment with tyrosine kinase inhibitors: review and recommendations for harmonizing current methodology for detecting BCR-ABL transcripts and kinase domain mutations and for expressing results. Blood. 108, 28–37 (2006).

12. S. Branford, L. Fletcher, N. C. Cross, M. C. Müller, A. Hochhaus, et al., Desirable performance characteristics for BCR-ABL measurement on an international reporting scale to allow consistent interpretation of individual patient response and comparison of response rates between clinical trials. Blood. 112, 3330–3338 (2008).

13. J. E. Cortes, C. Gambacorti-Passerini, M. W. Deininger, M. J. Mauro, C. Chuah et al., Bosutinib Versus Imatinib for Newly Diagnosed Chronic Myeloid Leukemia: Results From the Randomized BFORE Trial. J. Clin. Oncol. 36, 231–237 (2018).

14. M. Baccarani, M. W. Deininger, G. Rosti, A. Hochhaus, S. Soverini, et al., European LeukemiaNet recommendations for the management of chronic myeloid leukemia: 2013. Blood. 122, 872–884 (2013).

15. F. X. Mahon, G. Etienne. Deep molecular response in chronic myeloid leukemia: the new goal of therapy? Clin. Cancer Res. 20, 310–322 (2014).

16. D. M. Ross, I. S. Pagani, N. Shanmuganathan, C. H. Kok, J. F. Seymour, et al., Long-term treatment-free remission of chronic myeloid leukemia with falling levels of residual leukemic cells. Leukemia. 32, 2572–2579 (2018).

17. K. Sasaki, H. Kantarjian, S. O’Brien, F. Ravandi, M. Konopleva, et al., Prediction for sustained deep molecular response of BCR-ABL1 levels in patients with chronic myeloid leukemia in chronic phase. Cancer. 124, 1160–1168 (2018).

18. J. Hasford, M. Baccarani, V. Hoffmann, J. Guilhot, S. Saussele, et al., Predicting complete cytogenetic response and subsequent progression-free survival in 2060 patients with CML on imatinib treatment: the EUTOS score. Blood. 118, 686–692 (2011).

19. A. Hochhaus, M. Baccarani, R. T. Silver, C. Schiffer, J. F. Apperley, et al., European LeukemiaNet 2020 recommendations for treating chronic myeloid leukemia. Leukemia. 34, 966–984 (2020).

20. G. Clapp, D. Levy. A review of mathematical models for leukemia and lymphoma. Drug Discov. Today Dis. Models. 16, 1–6 (2014).

21. F. Michor, T. P. Hughes, Y. Iwasa, S. Branford, et al., Dynamics of chronic myeloid leukaemia. Nature. 435, 1267–1270 (2005).

22. M. Tang, M. Gonen, A. Quintas-Cardama, J. Cortes, H. Kantarjian, et al., Dynamics of chronic myeloid leukemia response to long-term targeted therapy reveal treatment effects on leukemic stem cells. Blood. 118, 1622–1631 (2011).

23. M. Horn, I. Glauche, M. C. Müller, R. Hehlmann, A. Hochhaus, et al., Model-based decision rules reduce the risk of molecular relapse after cessation of tyrosine kinase inhibitor therapy in chronic myeloid leukemia. Blood. 121, 378–384 (2013).

24. I. Roeder, M. Horn, I. Glauche, A. Hochhaus, M. C. Mueller, et al., Dynamic modeling of imatinib-treated chronic myeloid leukemia: functional insights and clinical implications. Nat. Med. 12, 1181–1184 (2006).

25. N. L. Komarova, D. Wodarz. Effect of cellular quiescence on the success of targeted CML therapy. PLoS One. 2, e990 (2007).

26. H. Ma, S. Leng, K. Aihara, W. Lin, L. Chen. Randomly distributed embedding making short-term high-dimensional data predictable. Proc. Natl. Acad. Sci. U.S.A. 115, E9994–E10002 (2018).

27. Y. Hirata, K. Morino, K. Akakura, C. S. Higano, N. Bruchovsky, et al., Intermittent Androgen Suppression: Estimating Parameters for Individual Patients Based on Initial PSA Data in Response to Androgen Deprivation Therapy. PLoS One. 10, e0130372 (2015).

28. K. Morino, Y. Hirata, R. Tomioka, H. Kashima, K. Yamanishi, et al., Predicting disease progression from short biomarker series using expert advice algorithm. Sci. Rep. 5, 8953 (2015).

29. S. Mustjoki, K. Auvinen, A. Kreutzman, P. Rousselot, S. Hernesniemi, et al., Rapid mobilization of cytotoxic lymphocytes induced by dasatinib therapy. Leukemia. 27, 914–924 (2013).

30. A. Hochhaus, T. Masszi, F. J. Giles, J. P. Radich, D. M. Ross. Treatment-free remission following frontline nilotinib in patients with chronic myeloid leukemia in chronic phase: results from the ENESTfreedom study. Leukemia. 31, 1525–1531 (2017).

31. M. Baccarani, J. Cortes, F. Pane, D. Niederwieser, G. Saglio, et al., Chronic myeloid leukemia: an update of concepts and management recommendations of European LeukemiaNet. J. Clin. Oncol. 27, 6041–6051 (2009).

