## Supplementary Information for "Early Dynamics of Chronic Myeloid Leukemia on Nilotinib Predicts Deep Molecular Response"

**This PDF file includes:**

Figures S1 to S5

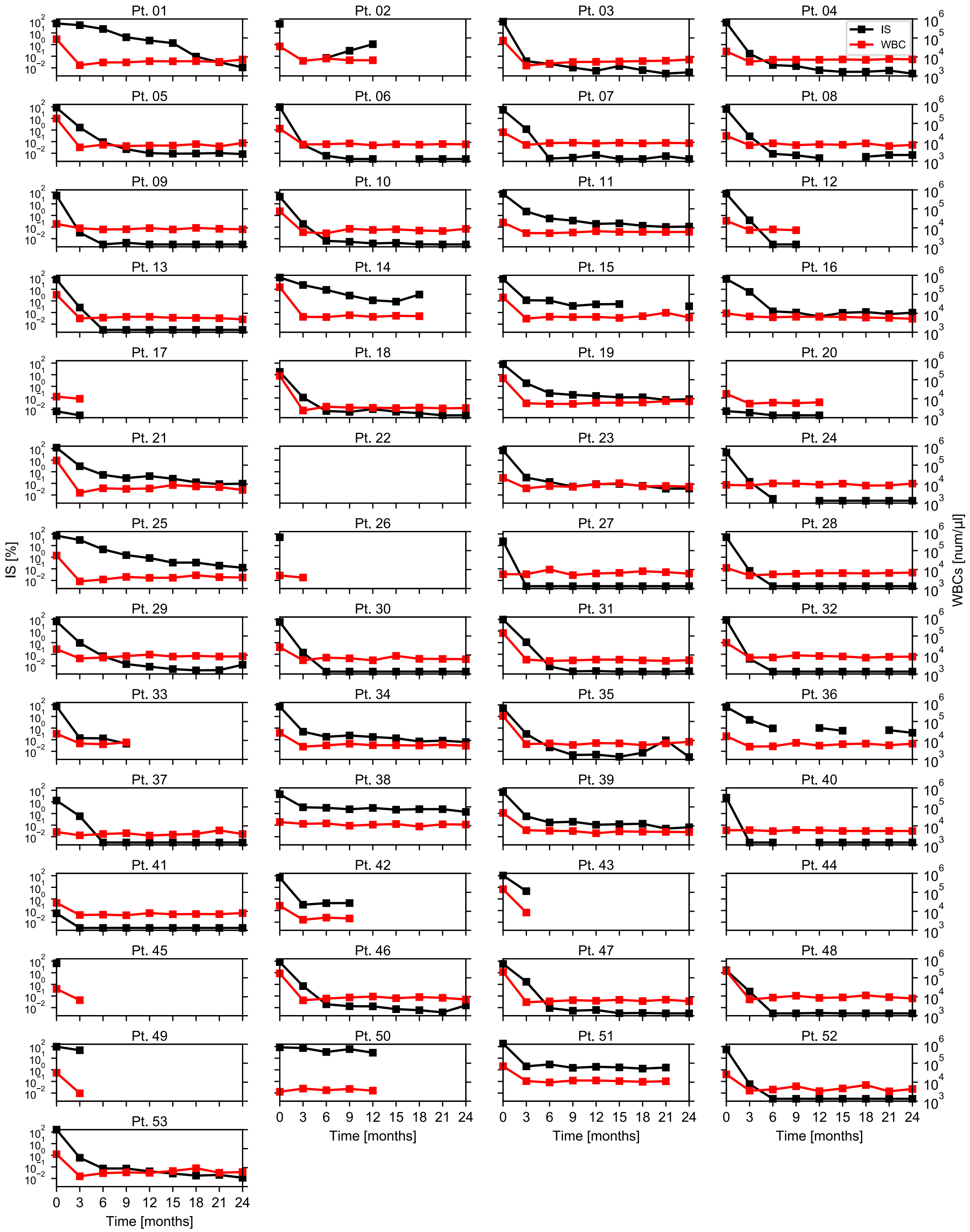

**Fig. S1. Measurement data for CML patients.**

Our prediction method used the time-series data for WBC counts and IS. The dataset consists of data from 53 patients. However, for 21 patients, the WBC count and/or IS data were incomplete at a certain time point. Thus, in this study, we used only the remaining 32 patient data.

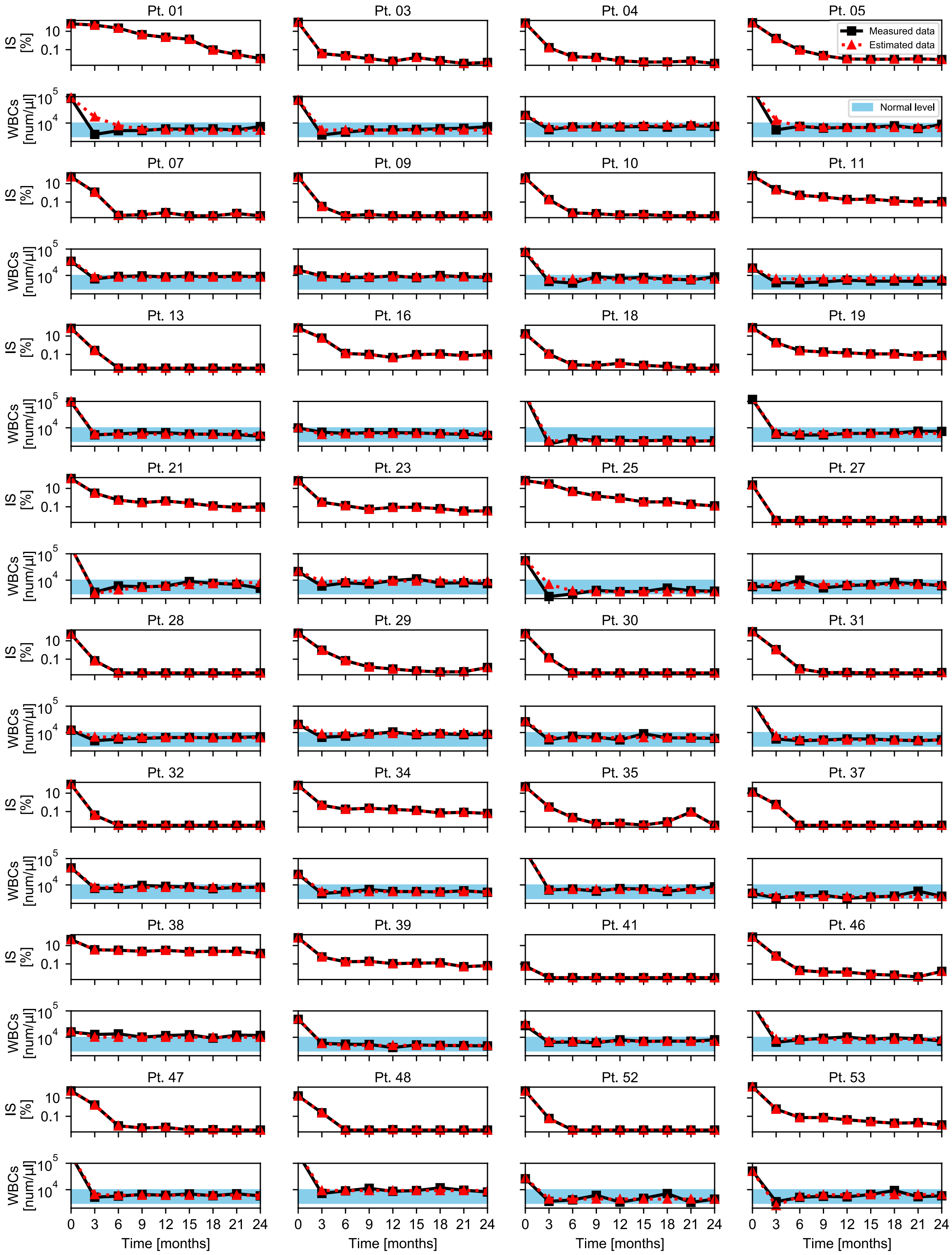


**Fig. S2. Our CML model approximated all measurement data.**
Each black solid line represents the WBC and IS measurement data for each patient. Each red dashed line represents the estimated dynamics based on our proposed model. Our model approximated all measurement data sufficiently.

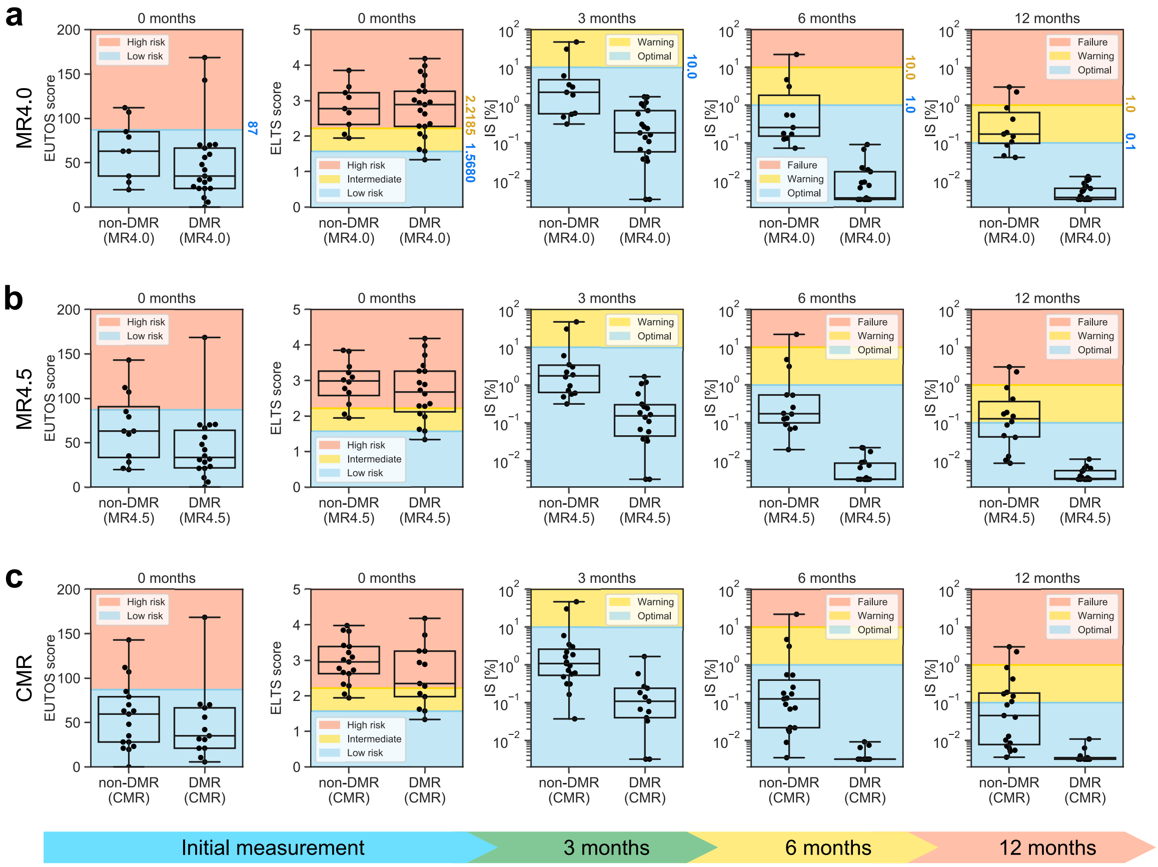


**Fig. S3. Effectiveness of the EUTOS score, ELTS score, and European LeukemiaNet guideline criteria for MR4.0, MR4.5, and CMR.**

**a-c,** Based on the approach described in Fig.1, we show the distribution of the EUTOS score, the ELTS score, and the ELN guideline for (a) MR4.0, (b) MR4.5, and (c) CMR. In most cases except for the guideline at 12 months for MR4.0 (the right top panel), these three approaches did not accurately predict non-MR4.0, non-MR4.5, and non-CMR patients.

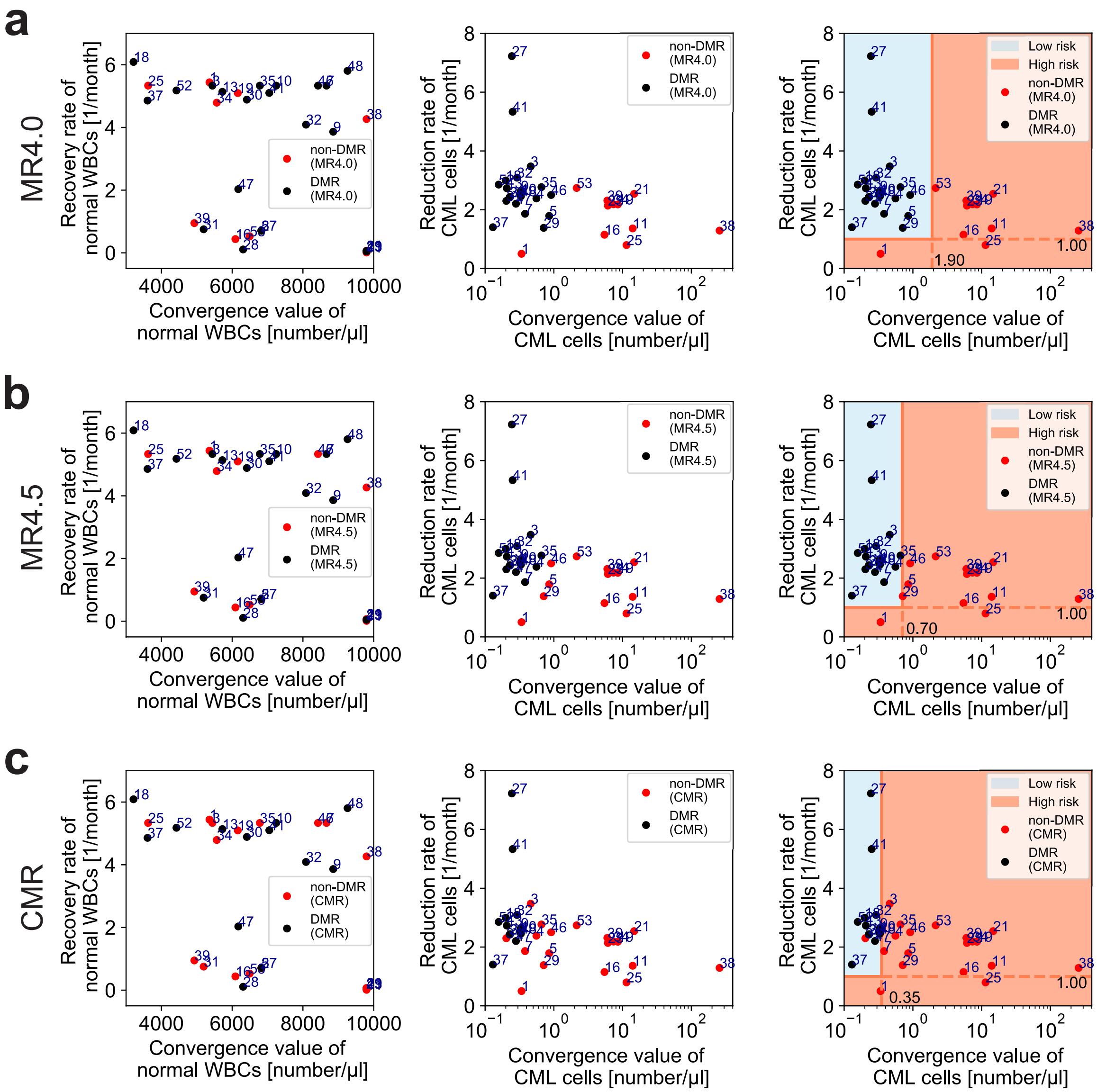


**Fig. S4. The classification of MR4.5 and CMR patients required the lower CML cell convergence value than that of MR4.0 patients.**

**a-c,** Distributions of the reduction rates and convergence values for (left) normal WBCs, (middle) CML cells, and (right) the classification. The labels of these figures are (a) MR4.0, (b) MR4.5, and (c) CMR.

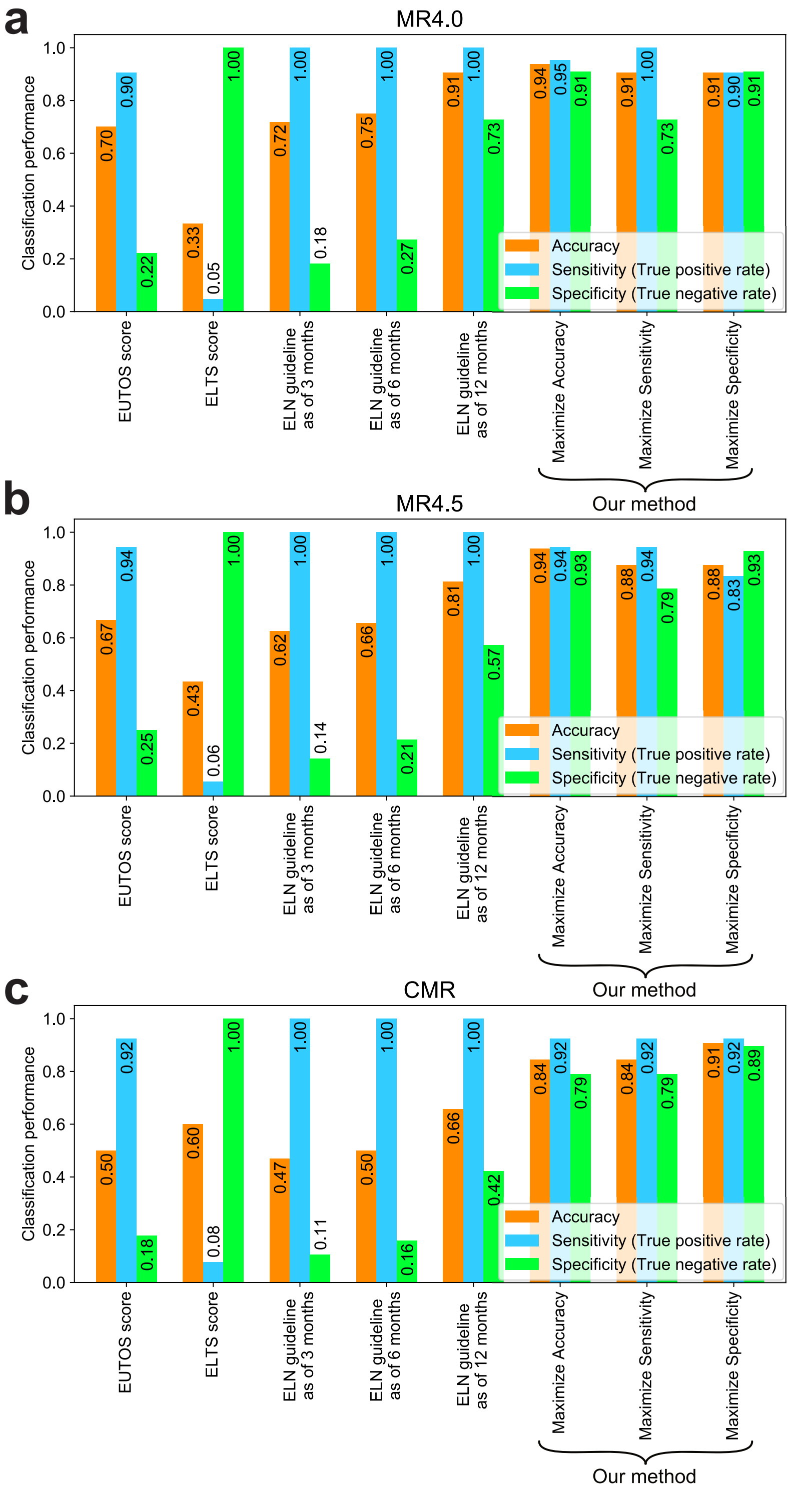


**Fig. S5. Our prediction method can also be applied to MR4.0, MR4.5 and CMR.**
**a-c**, Using the approach described in Fig. 4, we obtained the accuracy, sensitivity, and specificity performance for the EUTOS score, the ELTS score, the ELN guidelines, and our proposed method. The labels of these panels are (a) MR4.0, (b) MR4.5, and (c) CMR. The performance of our method preserved its performance even in CMR cases.
